# A clonal fresh water plants acquires transgenerational stress resistance under recurring copper excess

**DOI:** 10.1101/2021.02.06.430033

**Authors:** Meret Huber, Saskia Gablenz, Martin Höfer

## Abstract

Although non-genetic inheritance is thought to play an important role in plant ecology and evolution, evidence for adaptive transgenerational plasticity is scarce. Here, we investigated the consequences of copper excess on offspring defences and fitness in the giant duckweed (*Spirodela polyrhiza*) across multiple asexual generations. We found that exposing large monoclonal populations (>10,000 individuals) for 30 generations to copper excess decreased plant fitness during the first few generations but increased their fitness in consecutive generations under recurring stress when plants were grown for 5 generations under control conditions prior recurring conditions. Similarly, propagating individual plants as single descendants for 5 or 10 generations under copper excess decreased plant fitness when 5 generations and improved plant fitness when 10 generations passed between initial and recurring stress; thus, transgenerational stress responses likely contributed to the observed variations in offspring fitness of long-term copper exposed populations. Fitness benefits under recurring stress were partially associated with avoidance of excessive copper accumulation, which in turn correlated with transgenerationally modified flavonoid concentrations. Taken together, these data demonstrate time-dependent adaptive transgenerational responses under recurring stress, which highlights the importance of non-genetic inheritance for plant ecology and evolution.

## INTRODUCTION

Dwindling intraspecific genetic diversity in both natural and agroecosystems due to human practices has fuelled interest in the ability of species to resist environmental change in the absence of genetic variation through non-genetic inheritance. Non-genetic inheritance is any effect on offspring phenotype that is attributable to the transmission of factors other than the DNA sequence from ancestors [1], and includes vertical transmission of substances (e.g. nutrients, hormones, proteins, mRNA, toxins) [2, 3], epigenetic marks (DNA methylation, histone modification, non-coding small RNAs) [4, 5] and microbes [6]. Non-genetic inheritance may persist for one (parental), two (“multigenerational”) or more generations (“transgenerational”) [7, 8]. Theoretical work suggests that populations may acquire stress resistance through non-genetic inheritance, particularly when transmission fidelity across generations is high [9–11]. Experimental support of these predictions in multicellular organisms is however scarce [6, 12, 13], as such approaches require large-scale multi-generational studies and the ability to disentangle genetic from non-genetic factors.

In plants, a number studies showed that offspring may benefit from parental stress under recurring conditions (e.g. [4, 14–18]). To date, most of these studies focused on parental or multigenerational effects, and consequently, the observed patterns may partially be attributable to direct effects of the trigger on the organism during its early development [6, 19]. Furthermore, the adaptive value of transmitted traits is often inferred through indirect measurements of plant performance (but see [17, 20]), which may not adequately reflect fitness consequences [21]. Thus, directly assessing plant fitness when initial and recurring stress are separated by multiple generations is critical to progress our understanding in the ecological and evolutionary implications of non-genetic inheritance to mediate stress resistance.

Non-genetic inheritance might be particularly relevant in asexually reproducing plants [22], as this inheritance mode may compensate for the lack of genetic recombination [23, 24], may by-pass resetting during meiosis [19, 25, 26] and as the ancestor’s and offspring’s environments likely resemble during short-distance vegetative reproduction. Surprisingly however, research on transgenerational plasticity in clonally reproducing plants accumulated only relatively recently (e.g. [23, 24, 27–32]), despite the prevalence and importance of plant asexual reproduction for natural and agroecosystems [33–39]. In clonal plants, stress exposure may alter DNA methylation profiles across one or more generations [28, 31] and modulate offspring growth and traits [27–30]; however, transgenerational stability and fitness consequences of these responses remain mostly unclear. Furthermore, direct evidence that stress exposure benefits offspring fitness when multiple generations lie in between initial and recurring stress is lacking to date.

The giant duckweed, *Spirodela polyrhiza* (L.) Schleid., is a fresh water plant that produces rapidly and almost exclusively asexually through budding. This monocot often grows in proximity to agriculture and thus is often recurrently exposed to copper sulphate used in crop protection. Copper excess results in the formation of reactive oxygen species (ROS), which induce oxidative stress [40, 41].

Plants resistance to copper excess may involve either exclusion or neutralization of the metal ions [42–44]. Flavonoids, particularly *ortho*-dihydroxylated B-ring-substituted flavonoids, may help plants to resist copper excess through scavenging ROS and suppressing ROS-formation as chelating agents [45, 46]. *Spirodela polyrhiza* accumulates two *ortho*-dihydroxylated and two monohydroxylated B-ring-substituted flavonoids in high concentrations in its flat, thallus-like plant body (“fronds”) [47]. The *ortho*-dihydroxylated but not monohydroxylated B-ring substitute flavonoids are associated with copper resistance in *S. polyrhiza* [48].

Here, we investigated whether copper excess affects offspring defences and fitness in *S. polyrhiza* across generations. We found that descendants of large monoclonal *S. polyrhiza* populations that were exposed for 30 generations to copper excess had lower fitness under short and higher fitness during prolonged growth under recurring stress. These patterns resembled transgenerational stress responses, as propagating individual plants for 5 or 10 generations under copper excess had negative effects on plant fitness when 5 generations and positive effects when 10 generations passed between initial and recurring stress. Quantifying flavonoids and copper concentrations suggests that avoidance of excessive copper accumulation rather than enhanced induction of anti-oxidative flavonoids contributed to increased plant fitness under recurring stress.

## METHODS

### Plant material

*Spirodela polyrhiza* was cultivated under non-sterile conditions in N-medium (KH_2_PO_4_ 150 μM, Ca(NO_3_)_2_ 1 mM, KNO_3_ 8 mM, H_3_BO_3_ 5 μM, MnCl_2_ 13 μM, Na_2_MoO_4_ 0.4 μM, MgSO_4_ 1 mM, FeNaEDTA 25 μM in deionized water). Genotype 7498 from North Carolina (USA) was used for all experiments. Plants were grown in a climate chamber operating under the following conditions: 26°C constant; 18 h light; light intensity 130 – 160 μmol m^−2^ s^−1^, supplied by horizontally arranged neon tubes (Osram Lumilux L36 W/ 865 cool daylight; Osram GmbH, Munich, Germany).

### Statistical analysis

All statistical analysis was performed in R version 3.5.1 [49] using Rmsic [50], gridExtra [51], dplyr [52], ggplot2 [53], PerformanceAnalytics [54] and nlme [55]. More details on the statistical analyses are given in the experimental sections below.

### Long-term exposure experiment

In order to test whether *S. polyrhiza* acquires copper resistance in the absence of genetic variation, we grew replicated monoclonal *S. polyrhiza* populations in the presence and absence of copper excess for four months (approximately 30 generations; “pre-treatment”). Plants were grown inside 52 l transparent plastic ponds (79×58×17.5 cm, Bauhaus, Germany) that were filled with 30 l N-medium with or without 20 μM CuSO_4_ (N = 10 each). The ponds were covered with 4 mm transparent plexiglass (UV Gallery100, Sandrock, Germany) with 5 mm distance between pond edges and plates. Every two weeks – when the control populations covered approximately the entire pond – plants covering about 5% of the total pond surface were randomly chosen and transferred into refilled ponds. In the first four weeks of the experiment, the maximum population size per pond reached approximately 27,000 and 16,000 fronds in the control and copper medium, respectively.

Subsequently, algae colonized the ponds, which reduced maximum population size in the control but not copper medium to approximately 10,000.

Four months after start of the experiment, plant fitness of control and copper pre-treated populations was assessed in both environments across 16 days of growth (approx. 4-6 generations). To avoid growth bias due to direct effects of copper excess, plants were propagated for 5 consecutive generations as single descendants under control conditions. To this end, 5 fronds carrying a small daughter (generation 0) of each pond were transferred to transparent 50 ml polystyrene tubes (ø 2.8 cm, height 9.5 cm, Kisker, Germany) covered with foam plugs (Kisker, Germany) and filled with 30 ml control medium, with one frond per tube. The first daughter was separated from the mother and placed inside a new tube once the daughter had fully emerged.

Subsequently, to measure plant fitness in the absence of copper excess, a pool of the first daughter from generation four (5 fronds) of each pond was placed inside one half of the 18 L containers that were divided in the middle by a fine mesh, filled with 10 l N-medium and covered with transparent PET lids (Pöppelmann, Lohne, Germany). Each container received the fronds of copper and control pre-treated populations to avoid growth bias due to potentially co-evolved microbes. To measure plant fitness in the presence of copper excess, a pool of the second daughters from generation four of each pond were placed inside containers filled with N-medium containing 20 μM CuSO_4_ as described above. After 8 days of growth, the total number of fronds was counted and the growth medium exchanged. After 16 days, the number of fronds was counted once more, plants were subsequently dried with a tissue paper and fresh weight was determined. Relative growth rates (RGR) were calculated as RGR = (ln(N_2_) – ln(N_1_))/(t_2_-t_1_) [56], with N = population size and t = day. Pre-treatment effects on plant fitness were assessed by comparing RGR between pre-treatments for each offspring environment separately using Kruskal-Wallis rank sum tests. Pre-treatment effects on RGR were expressed as the ratio in RGR of copper to control pre-treated offspring. Resistance was analysed as the difference in these pre-treatment effects on RGR between offspring environments using Kruskal-Wallis rank sum tests. To test for interactions of the pre-treatment and growth interval on RGR, two-way analysis of variance (ANOVAs) were performed. To test for interactions of the pre-treatment and offspring environment on RGR and biomass (log) accumulation, two-way ANOVAs were performed. Biomass accumulation was log-transformed (natural logarithm) prior statistical analysis to account for the exponential growth of *S. polyrhiza*.

### Transgenerational experiment

To investigate transgenerational stress responses, we propagated individual *S. polyrhiza* plants as single descendants for different durations under copper and control conditions (Fig S1 in Supplemental Information). Individual mother plants carrying a small daughter were placed inside 50 ml polystyrene tubes filled with 30 ml N-medium containing or lacking 20 μM CuSO_4_, and grown as single descendants as described above for 2, 5 or 10 generations (“pre-treatment”; N = 25 each). The starting time points of the control lineages were delayed to allow simultaneous assessment of plant fitness later on. After the pre-treatment phase, propagation continued for 5, 10 or 15 generations in control medium (“recovery”), after which offspring growth assays inside transparent 250 ml polypropylene beakers (Plastikbecher GmbH, Giengen an der Brenz, Germany) filled with 180 ml medium and covered with a perforated and transparent lid were performed. For these assays, the first daughters were transferred to copper-free medium, whereas the second daughters were subjected to medium containing 20 μM CuSO_4_ (“initial fronds”). An image for surface area analysis was taken at a camera installation with a webcam (HD Pro Webcam C920, Logitech; webcam software Yawcam version 0.6.0) and a subjacent adjustable LED light after setting up the experiment. During the growth assays, offspring that directly emerged from the initial fronds were marked (“direct offspring”). The growth medium was exchanged four days after the start of the experiment. After 8 days of growth, the number of direct offspring was counted, an image was taken, fronds were gently dried with a tissue paper and the fresh weight of the initial frond and all other plants (“offspring”) was determined. The initial frond and the offspring were flash-frozen separately inside tubes in liquid nitrogen and stored at −80 °C until further analysis.

Frond surface area was measured with ImageJ (version 2.0.0-rc-43/1.51k; Java version 1.6.0_24). Flavonoid concentrations were measured on an HPLC 1100 series equipment (Agilent Technologies) coupled to a photodiode array detector (G1315A DAD, Agilent Technologies) and quantified as described in [48]. To measure plant copper concentration, 20-50 mg ground plant material was extracted with 1 ml 0.01 M MOPS buffer. Samples were centrifuged at 17,000 g and supernatant diluted 1:2 and 1:3 in 0.01 M MOPS buffer for control and copper-subjected plants, respectively. Copper concentration was measured using a Copper Assay Kit (Sigma Aldrich) following the manufacturer’s instructions on an Infinite^®^ 200 PRO NanoQuant Plate Reader Spectrophotometers (Tecan Trading AG, Männedorf, Switzerland) at 359 nm. Absolute concentrations were calculated based on an external standard curve.

Differences in biomass (log) accumulation, surface area, area- and biomass-based growth rates, the fresh weight of the initial frond and the number of offspring of the initial frond between copper and control pre-treated offspring were analyzed with Kruskal-Wallis rank sum tests for each assay and offspring environment separately. To test for interactions of the pre-treatment and the offspring environment on these parameters, two-way ANOVAs were performed. Growth rates were calculated as described above with N = surface area or fresh weight, respectively. Fitness effect of copper pre-treatment was expressed as the ratio in biomass (log) accumulation of copper to control pre-treated offspring. Resistance was analyzed as the difference in these pre-treatment effects between offspring environments using Kruskal-Wallis rank sum tests. Differences in flavonoid concentrations and plant copper concentration between copper and control pre-treated offspring were analyzed with Student’s *t*-tests for each assay and offspring environment separately. To test for interactions of the pre-treatment and offspring environment on flavonoid and copper accumulation, two-way ANOVAs were performed. Pre-treatment effects on flavonoid and copper accumulation were expressed as the ratio in flavonoid or copper concentration of copper to control pre-treated offspring, respectively. The sum of the two major mono-(apigenin 8-C- and 7-O-glucoside) and dihydroxylated (luteolin 8-C- and 7-O-glucoside) B-ring substituted flavonoids were analyzed separately. Due to experimental errors, the number of replicates was reduced in some assays. The correlation among the different fitness parameters, i.e. offspring biomass (log) accumulation, the number of direct offspring and initial frond fresh weight, surface area (log), surface-area and biomass-based growth rates, were calculated using Pearson’s moment correlations. Mixed effect models were applied to assess the correlation between pre-treatment effects on offspring copper and flavonoid concentrations, as well as between offspring copper and biomass (log) accumulation using maximum likelihood estimations with assay as a random effect. Significant correlations were assessed by comparing two models with and without flavonoid or biomass (log) accumulation, respectively, as a fixed effect using ANOVA. For the correlation between pre-treatment effects on copper and biomass (log) accumulation, additional models were analyzed in which the data point with extremely high pre-treatment effect on biomass accumulation (> 1.4) was excluded.

## RESULTS

To test whether *S. polyrhiza* may acquire stress resistance in the absence of genetic variation, we grew large monoclonal populations for 30 generations in the presence and absence of copper excess (“pre-treatment”) and subsequently propagated offspring for 5 generations under control conditions prior measuring offspring growth rates across 16 days. In the first 8 days of growth, copper pre-treatment reduced offspring growth rates under both control and copper excess to a similar extent (−16%, *P* < 0.001 and −26%, *P* = 0.04, respectively, Kruskal-Wallis rank sum tests; Fig 1 and Fig S2); thus, the pre-treatment did not alter plant resistance (difference in the pre-treatment effects on plant growth rates between offspring environment; *P* = 0.20, Kruskal-Wallis rank sum tests, Fig 1). In the consecutive 8 days, copper pre-treatment enhanced offspring growth rates under copper excess (+36%, *P* = 0.005) while it had no effect under control conditions (*P* = 0.56, Kruskal-Wallis rank sum tests, Fig 1, Fig S2); consequently, plant resistance increased upon copper pre-treatment (*P* = 0.003, Kruskal-Wallis rank sum test, Fig 1). Across the entire 16 days, copper pre-treatment reduced growth rates under control conditions (−14%, *P* = 0.002) and had no effect on growth rates under copper excess (*P* = 0.94, Kruskal-Wallis rank sum tests, Fig S2). Consequently, copper pre-treatment reduced growth depression by copper excess across these 16 days of growth (*P* < 0.001, two-way ANOVA, Fig S2). Assessing plant resistance and fitness based on plant biomass accumulation instead of growth rates exhibited similar patterns (Fig S3). Taken together, these data show that long-term growth of *S. polyrhiza* under copper excess may benefit plant fitness and resistance under recurring stress.

**Figure 1.**
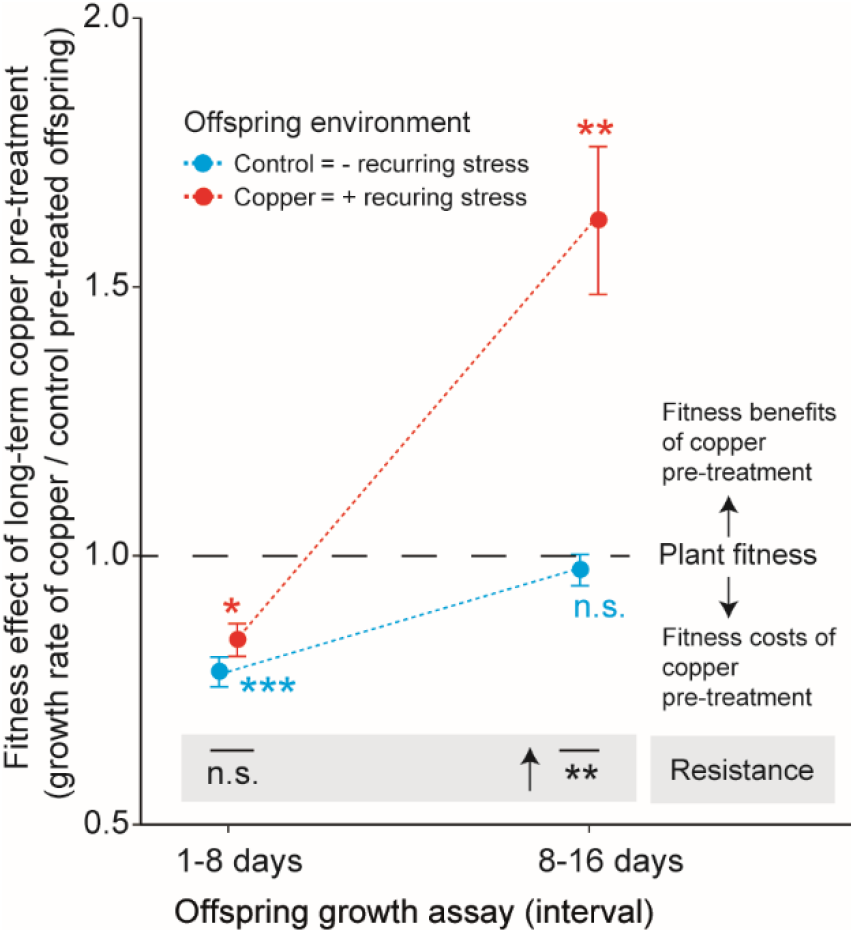
Growth of large monoclonal *S. polyrhi*z*a* populations for 30 generations under copper excess had negative effects on offspring fitness during short and positive effects during prolonged growth under recurring copper excess. Fitness effects of long-term copper pre-treatment are the ratios in growth rates of copper to control pre-treated offspring. Asterisks beside data points depict *P*-values of Kruskal-Wallis rank sum tests comparing growth rates of copper and control pre-treated offspring for each offspring environment and growth interval separately. Resistance is the difference in fitness pre-treatment effects between offspring environments (Kruskal-Wallis rank sum tests, arrow indicates positive effect). Descendants from copper and control pre-treated populations were grown for 5 generations under control conditions prior assessing offspring growth. (\**P* < 0.05, \*\**P* < 0.01, \*\*\**P* < 0.001; n.s. = non-significant). Data display mean values and standard errors. N = 9-10.

The observed variation in plant fitness and resistance of long-term pre-treated populations may have been the consequence of selection of newly acquired phenotypes or due to transgenerational stress responses in the absence of selection. To investigate on the latter, we investigated offspring defence and fitness after different durations under copper excess (“pre-treatment”; 2-10 generations) as well as after different durations under control conditions prior recurring stress (“recovery”; 5-15 generations) using single descendant propagations, in which no selection takes place (Fig S1). Under 5 generations recovery, two generations of copper pre-treatment had positive effects on plant biomass accumulation both in the presence as well as in the absence of recurring stress. Prolonging the pre-treatment to 5 generations eliminated these beneficial effects in both offspring environment. Further expanding the pre-treatment to 10 generations resulted in negative effects on biomass accumulation in the presence of recurring stress, and neutral effects in the absence of recurring stress (Kruskal-Wallis rank sum tests, Fig 2, Fig S4). Thus, under 5 generations recovery, copper pre-treatment had neutral (two and 5 generations pre-treatment) or negative effects (10 generations pre-treatment) on plant resistance (Kruskal-Wallis rank sum tests, Fig 2). Under 10 generations recovery, two generation pre-treatment elicited positive effects on plant biomass accumulation in the presence and absence of recurring stress, similar to the observations under 5 generations recovery. However, prolonging the pre-treatment to 5 and 10 generations had beneficial effects on offspring biomass accumulation in the presence and negative effects in the absence of recurring stress (Kruskal-Wallis rank sum tests, Fig 2, Fig S4). Thus, plant resistance increased under these pre-treatment and recovery combinations (Kruskal-Wallis rank sum tests, Fig 2). Under 15 generations recovery, pre-treatment did not affect biomass accumulation in the presence or absence of stress (Kruskal-Wallis rank sum test, *P* = 0.94, Fig 2, Fig S4). Other fitness parameters, i.e. surface area, growth rates based on surface area and biomass accumulation, as well as the initial plant’s fresh weight and offspring number, were all strongly correlated with each other and with offspring biomass accumulation (R^2^ > 0.49, *P* < 0.001, Pearson’s correlation tests, Fig S5), and exhibited similar pre-treatment effects (Figs S6-S10), thus corroborating the above described findings based on offspring biomass accumulation. Taken together, these data demonstrated that copper excess may affect plant fitness and resistance under recurring conditions, and the magnitude and direction of the effects are dependent on the duration of the pre-treatment and recovery phase.

**Figure 2.**
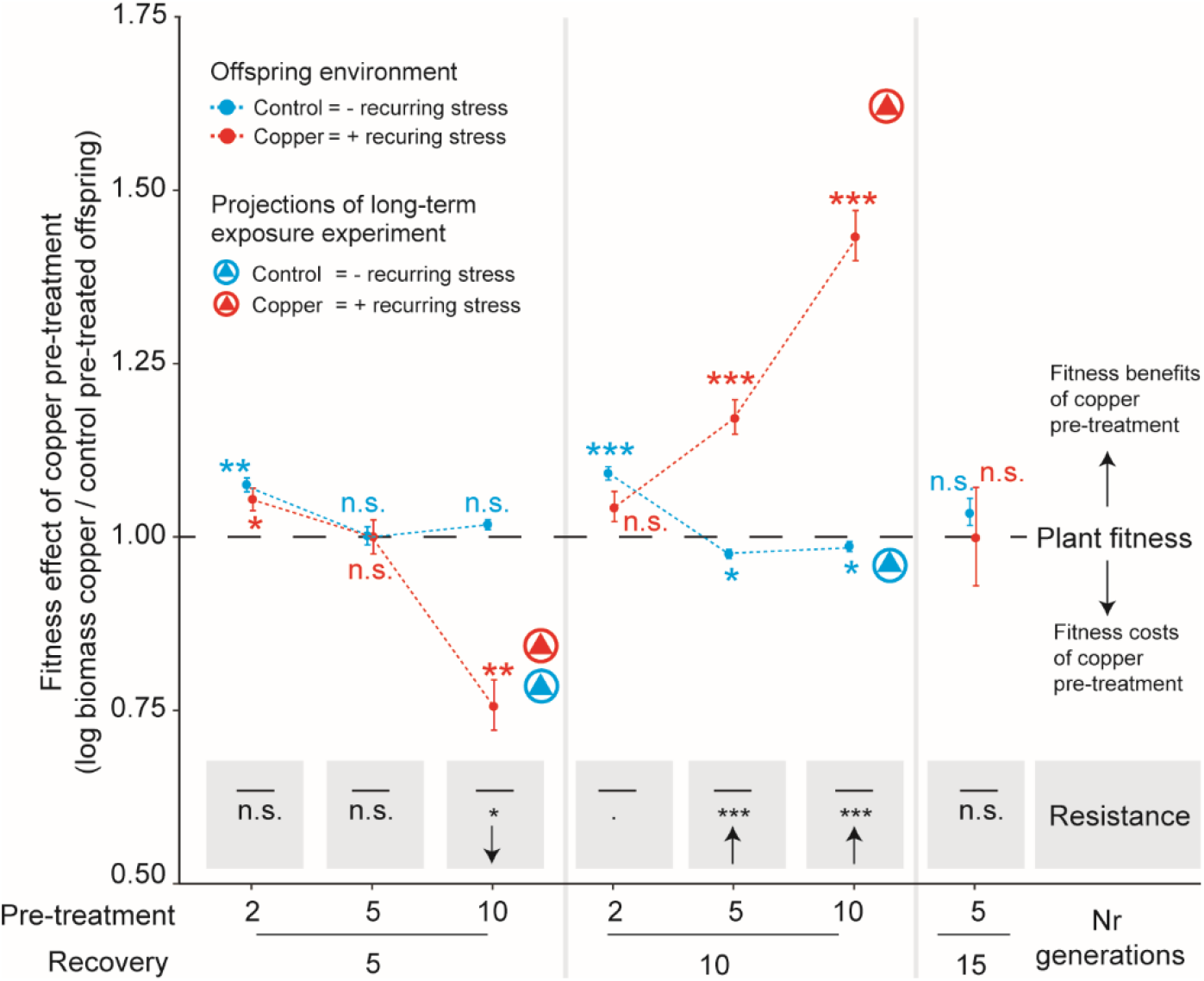
Copper pre-treatment under single descendant propagation can elicit positive fitness and resistance effects under recurring stress, similar to pre-treatment effects observed after long-term growth of monoclonal *S. polyrhiza* populations. Fitness effects of copper pre-treatment are the ratios of biomass (log) accumulation of copper to control pre-treated offspring after 8 days of growth for both offspring environments separately. Asterisks next to data points indicate *P*-values of Kruskal-Wallis rank sum tests comparing biomass (log) accumulation of copper and control pre-treated offspring within each offspring environment. Resistance is the difference in these pre-treatment effects between offspring environments (Kruskal-Wallis rank sum tests, arrows indicate positive and negative effects). Mean values of the long-term copper exposure are projected as encircled triangles. (\**P* < 0.05, \*\**P* < 0.01, \*\*\**P* < 0.001; n.s. = non-significant). Data display mean values and standard errors. N = 3-20.

To assess whether pre-treatment effects on biomass accumulation were due to altered accumulation of defensive metabolites, we measured the concentrations of the four major flavonoids in offspring. Depending on the pre-treatment and recovery phase combination, copper pre-treatment enhanced the accumulation of the dihydroxylated B-ring substituted flavonoids, including elevated basal and induced levels as well as only elevated induced levels (= primed) after up to 15 generations of recovery (Student’s *t*-tests, Fig 3). However, in the assays that exhibited transgenerationally elevated flavonoid levels, no alteration in plant resistance by copper pre-treatment was observed (Fig 3). In contrast, in the two pre-treatment and recovery phase combinations with increased plant fitness and resistance under recurring stress (5 and 10 generations pre-treatment, 10 generations recovery), copper pre-treatment had either neutral (10 generations recovery) or negative (5 generations recovery) effects on the concentration of the dihydroxylated B-ring substituted flavonoids under control conditions, and no effect on the accumulation of these metabolites under recurring stress (Student’s *t*-tests, Fig 3). Across all assays, the pre-treatment elicited very similar effects on all four major flavonoids (Figs S11-S15) - albeit with stronger effects in the di – than monohydroxylated B-ring substituted flavonoids - except that the monohydroxylated B-ring substituted flavonoids were not primed (*P* = 0.7, Student’s *t*-tests, Fig S13). Taken together, these data show that copper pre-treatment may transgenerationally increase, decrease and prime dihydroxylated B-ring substituted flavonoids when initial and recurring stress were separated by up to 15 generations, and indicate that flavonoids may not be the sole contributor for enhanced offspring fitness under recurring stress.

**Figure 3.**
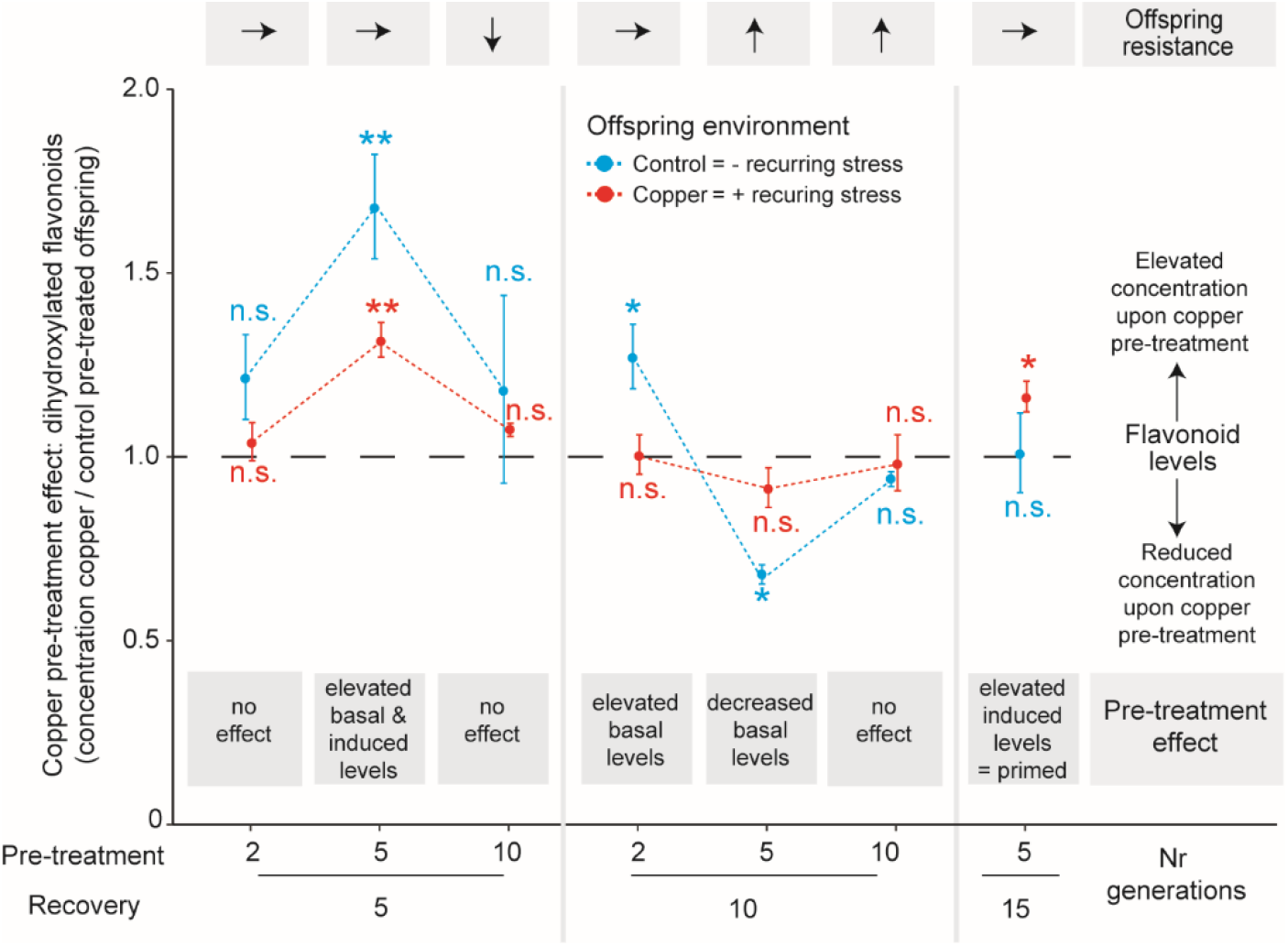
Copper pre-treatment may enhance offspring flavonoid accumulation, but these increments were not associated with enhanced offspring resistance. Copper pre-treatment effects on di-hydroxylated B-ring substituted flavonoid accumulation were expressed as the ratio in the sum of the two major luteolin glucosides of copper to control pre-treated offspring after 8 days of growth, for both offspring environments separately. Asterisks display *P*-values of Student’s *t*-tests comparing dihydroxylated B-ring substituted flavonoid levels of copper and control pre-treated offspring within each offspring environment. Offspring resistance is display above the panel as arrows and refer to results of figure 2 (\**P* < 0.05, \*\**P* < 0.01, \*\*\**P* < 0.001; n.s. = non-significant). Data display mean values and standard errors. N = 3-6.

To investigate whether altered copper uptake or excretion efficiency may account for the observed pre-treatment effects in plant fitness and flavonoid accumulation, we measured offspring copper concentration in a subset of these assays (Fig 4, Fig S16). In the assay in which copper pre-treatment benefited plant resistance and reduced basal flavonoid concentrations (5 and 10 generation exposure and recovery, respectively), offspring copper concentrations were lower in copper compared to control pre-treated offspring (pre-treatment: *P* = 0.06, two-way ANOVA), particularly in the presence of recurring stress (*P* = 0.049, Student’s *t*-test, Fig 4). In the second assay with beneficial effects of copper pre-treatment on offspring resistance (10 generations pre-treatment and recovery each), pre-treatment did not affect plant copper accumulation (Student’s *t*-tests, *P* > 0.86, Fig S16). In the assay with elevated basal and induced flavonoid levels (5 generations exposure and recovery each), copper pre-treated offspring exhibited increased copper concentration (*P* = 0.009, two-way ANOVA), both in the presence (*P* = 0.09) and absence (*P* = 0.07, Student’s *t*-tests) of recurring stress compared to control pre-treated offspring (Fig S16). Across all assays, pre-treatment effects of copper accumulation tended to be negatively correlated with pre-treatment effects of offspring biomass (log) accumulation (*P* = 0.06, mixed effect models, Fig 5A), but only if the assay with extremely high benefits of copper pre-treatment on plant fitness under recurring stress was excluded from the analysis. Furthermore, pre-treatment effects of copper accumulation were positively correlated with pre-treatment effects of total and individual flavonoid accumulation (*P* = 0.02; mixed effect models, Fig 5B, Fig S17). Thus, pre-treatment effects of plant copper accumulation closely reflected pre-treatment effects of plant flavonoid concentrations, and avoidance of excessive copper accumulation in copper pre-treated offspring was partially associated with improved plant fitness and resistance under recurring stress.

**Figure 4.**
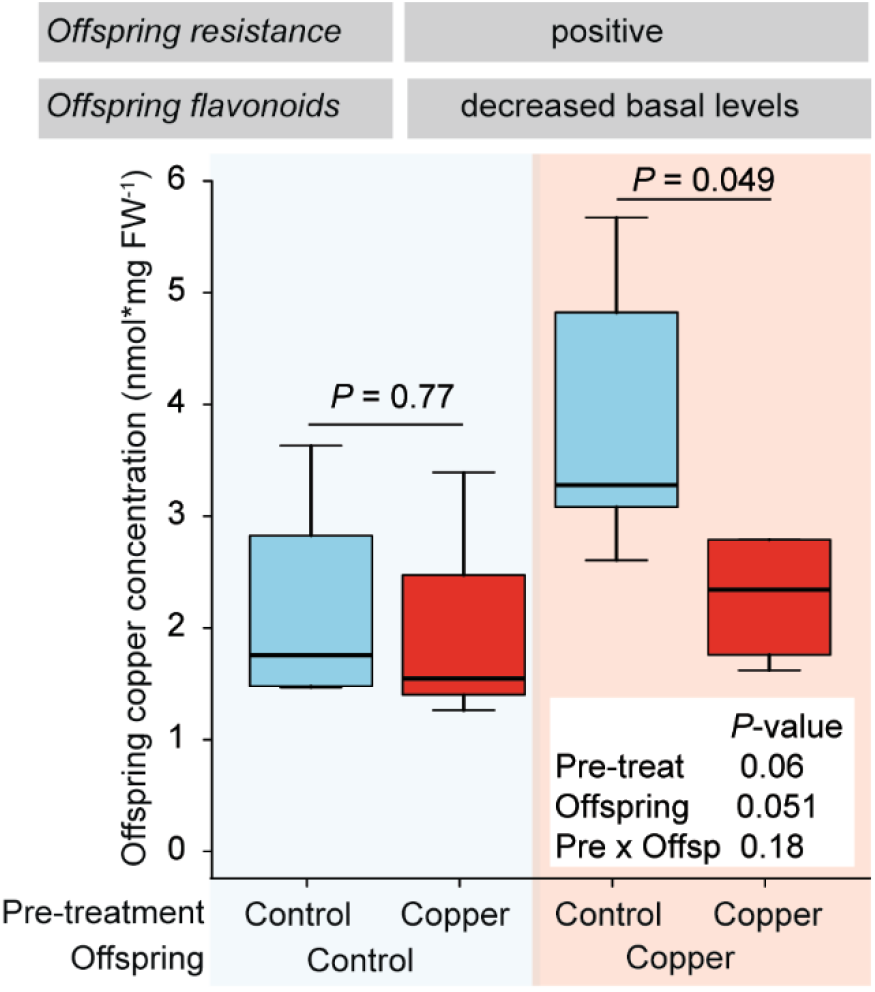
Avoidance of excessive copper accumulation was partially associated with enhanced offspring resistance under recurring stress. Offspring copper concentration was measured after 8 days of growth in the presence and absence of copper excess in the assay with 5 and 10 generations of pre-treatment and recovery, respectively. *P*-values of Student’s *t*-tests and two-way ANOVAs are shown. Offspring resistance and flavonoid levels are display above the panel and refer to results of figure 2 and 3. N = 4-5.

**Figure 5.**
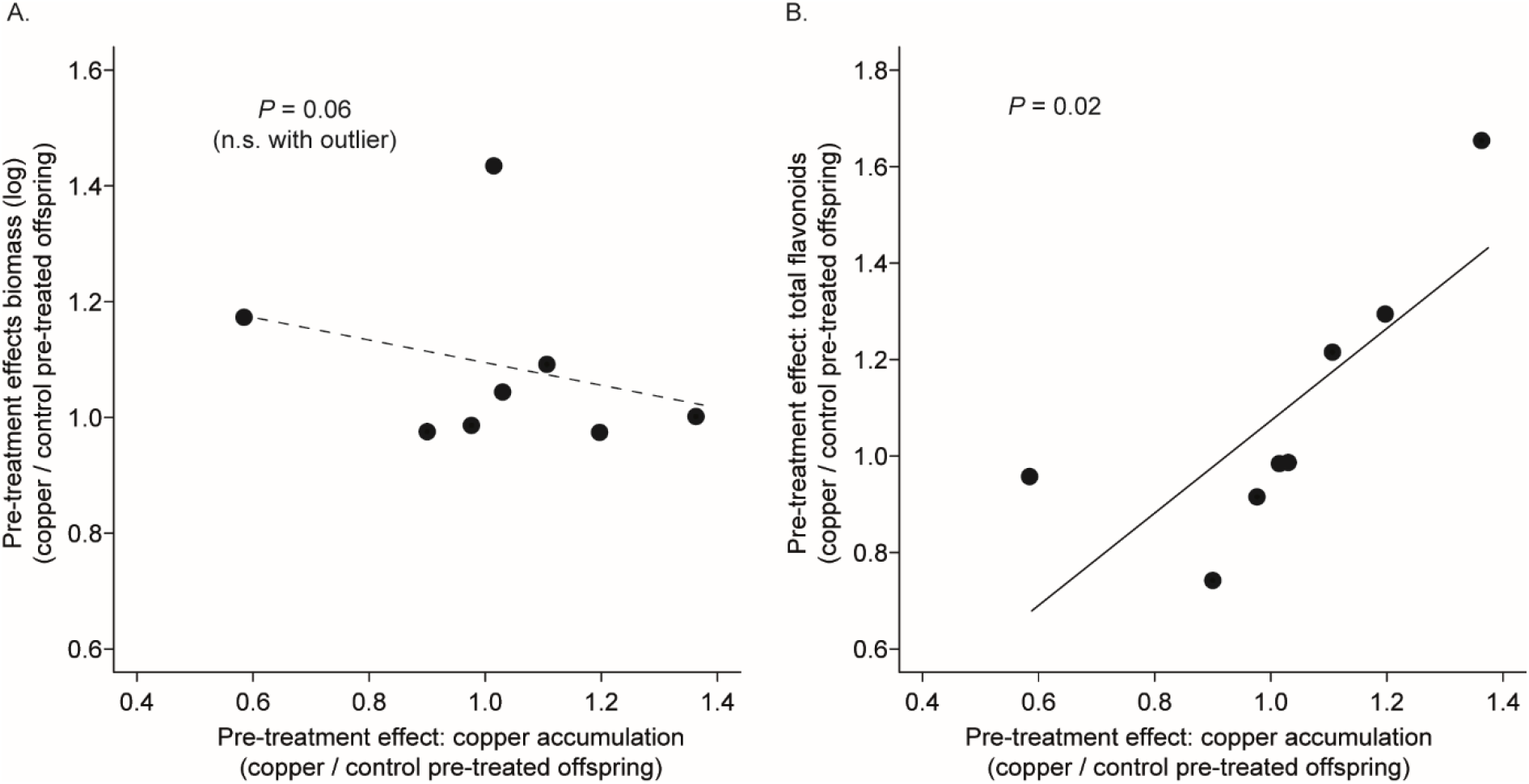
Pre-treatment effects of copper accumulation correlated weakly negatively with pre-treatment effects of biomass accumulation and positively with pre-treatment effects of flavonoid concentrations. Pre-treatment effects are the ratio in copper concentration and offspring biomass accumulation (log) (A), as well as total flavonoids concentration (B), of copper to control pre-treated offspring. Each data point displays the mean value of a pre-treatment and recovery combination and offspring environment. *P-*values of the comparison of linear mixed effect models using one-way ANOVA are shown. Results of models including and excluding the extreme pre-treatment effect in biomass accumulation (> 1.4) are shown in (B). N = 7-8. n.s. = not significant.

## DISCUSSION

The importance of non-genetic inheritance is a major controversy in plant ecology and evolution [8, 26, 57–59], also because clear evidence for adaptive responses across multiple generations is scarce. Here, we showed that in the clonal fresh water plant *S. polyrhiza* copper excess may have positive effects on offspring fitness when initial and recurring stress conditions were separated by multiple generations. Thereby, this study underlines the notion that non-genetic inheritance plays an important role in the ecology and evolution of asexual plants.

When large monoclonal populations of *S. polyrhiza* were grown for four months (approximately 30 generations) under copper excess, offspring of copper pre-treated populations exhibited under recurring stress lower plant fitness in the first 8 days of growth (2-3 generations) and higher plant fitness in the consecutive 8 days compared to offspring of control pre-treated populations, even though plants were grown for 5 generations in the absence of stress prior fitness assays. Genetic mutations arise very slowly in *S. polyrhiza* (approximately 0.004 point mutation in the protein coding sequence per generation) [60] and thus very unlikely account for the observed patterns. Instead, these variations in fitness may on the one hand be the result of intra-clonal selection of new phenotypes that were either induced by copper excess or randomly occurred trough non-genetic processes [61, 62]. On the other hand, such patterns may be due to transgenerational responses, in which the pre-treatment alters offspring fitness without selection of phenotypic variants [11, 17, 20, 26, 58]. Indeed, variation in fitness between long-term copper and control pre-treated populations were similar to the observed transgenerational responses during single descendant propagation. Thus, transgenerational responses likely contributed to the observed variation in plant fitness between long-term copper and control pre-treated populations.

In our transgenerational experiment, we found that copper pre-treatment may have positive effects on offspring fitness in the presence and negative effects in the absence of recurring stress when 10 generations separated initial and recurring stress. To our knowledge, such long-term transgenerational effects on plants fitness have not been reported to date. In most studies, transgenerational effects on plant performance and phenotype vanish after one to two generations [4, 11, 58, 63, 64], but see [28–30, 65, 66]. As most studies however use indirect estimates of plant fitness such as defensive phenotypes and biomass accumulation in sexual plants, which may not adequately reflect fitness consequences [6, 21], the adaptive value of the observed responses are often equivocal. By assessing plant fitness based on biomass accumulation in an asexual, thallus-like plant, we could directly demonstrate that copper excess may have both positive as well as negative effects on offspring fitness when multiple generations separate initial and recurring stress.

Interestingly, we observed that copper pre-treatment can have negative plant fitness effects after short (5 generations), and positive effects after long (10 generations) recovery period. Similarly, offspring flavonoid concentrations showed a large variation of different pre-treatment patterns including elevated basal and induced levels, as well as priming, and these effects did not simply weaken with increasing recovery time. Such contrasting effects in plant phenotypes and fitness across generations have been reported previously in both plants and animals [59, 67]. This shows that transgenerational effects may not simply decay over generations, as often predicted and observed [11, 65, 67–69], and highlights the importance of the timing in transgenerational responses [68].

Our analysis on offspring phenotypes suggest that reduced copper accumulation under recurring stress may account for beneficial effects of copper pre-treatment on offspring fitness under recurring stress. First, in the only assay in which copper pre-treatment reduced offspring copper concentration, copper pre-treatment benefited offspring fitness under recurring stress. Second, transgenerational effects of copper and biomass accumulation tended to be negatively correlated. Avoidance of excessive metal ion accumulation is a common strategy to resist toxic heavy metal concentrations [42, 43]. In contrast, no association between transgenerational effects on plant fitness and flavonoids were observed, despite that flavonoids, particularly the di-hydroxylated B-ring substituted compounds, are thought to be defensive and are associated with heavy metal resistance in *S. polyrhiza* and other plants [46, 48, 70]. Instead, pre-treatment effects of flavonoid and copper accumulation were positively correlated, suggesting that different copper accumulation triggered transgenerational patterns of flavonoid accumulation. Experiments that manipulate flavonoid concentrations as well as plant copper accumulation may provide further mechanistic insights into the relative importance of these physiological responses in modulating plant fitness across generations.

While we started to uncover the physiological mechanisms that mediate benefits of copper pre-treatment for offspring fitness under recurring stress, the underlying molecular basis remains unknown. Vertical transmission of substances very unlikely caused the observed pre-treatment effects, as any substance will have reached neglectable concentrations after the multigenerational recovery phase. In contrast, alterations of the microbial community through toxic copper concentrations may have contributed to the observed pre-treatment effects, particularly in our transgenerational experiment, which was performed under non-sterile conditions and without exchange of microbial communities among treatment groups. Transgenerational effects that were mediated by an altered microbial community have been reported in antibiotic-treated *Drosophila melanogaster* larvae [71]. In our long-term pre-treatment experiment, offspring of copper and control pre-treated populations were grown together; the observed fitness difference between copper and control pre-treated offspring are thus unlikely due to alterations in the mobile microbial community. Considering that many of the observed variations between copper and control pre-treated offspring were still detected after 10 (plant fitness) and even 15 (priming of flavonoids) generations, an involvement of epigenetic inheritance seems likely. Experiments that assess induction and persistence of epigenetic marks upon copper excess and their relation to plant phenotype and fitness may help to resolve the on-going controversy about the importance of epigenetic inheritance to mediate transgenerational stress resistance [5, 8, 13, 26, 57].

Taken together, our data demonstrate time-dependent costs and benefits of copper pre-treatment on offspring fitness when initial and recurring stress conditions were separated by multiple generations. Thereby, this study supports the notion that non-genetic inheritance may modulate plant ecology and evolution in asexually reproducing plants, which may help to explain the ecological and evolutionary success of these organisms.

## ACKNOWLEDGEMENTS

We thank Daniel Veit for crafting experimental containers, Michael Reichelt for supporting HPLC analysis and Jonathan Gershenzon for financial support, and Shuqing Xu, Martin Schäfer, Martin Höfer, Jonathan Gershenzon and Alexandra Chavez for helpful comments on the manuscript. This work was funded by the Max-Planck Society, the University of Münster and the Swiss National Science foundation (grant P400PB_186770 to Meret Huber). This paper is dedicated to our friend and colleague Saskia Gablenz.

## Author contributions

Conceived and designed the experiments: MeH. Performed the experiments: SG, MaH. Analyzed data: MeH, SG, MaH. Wrote manuscript: MeH.

