## Supplemental Information for "A clonal fresh water plants acquires transgenerational stress resistance under recurring copper excess"

SUPPLEMENTAL FIGURES

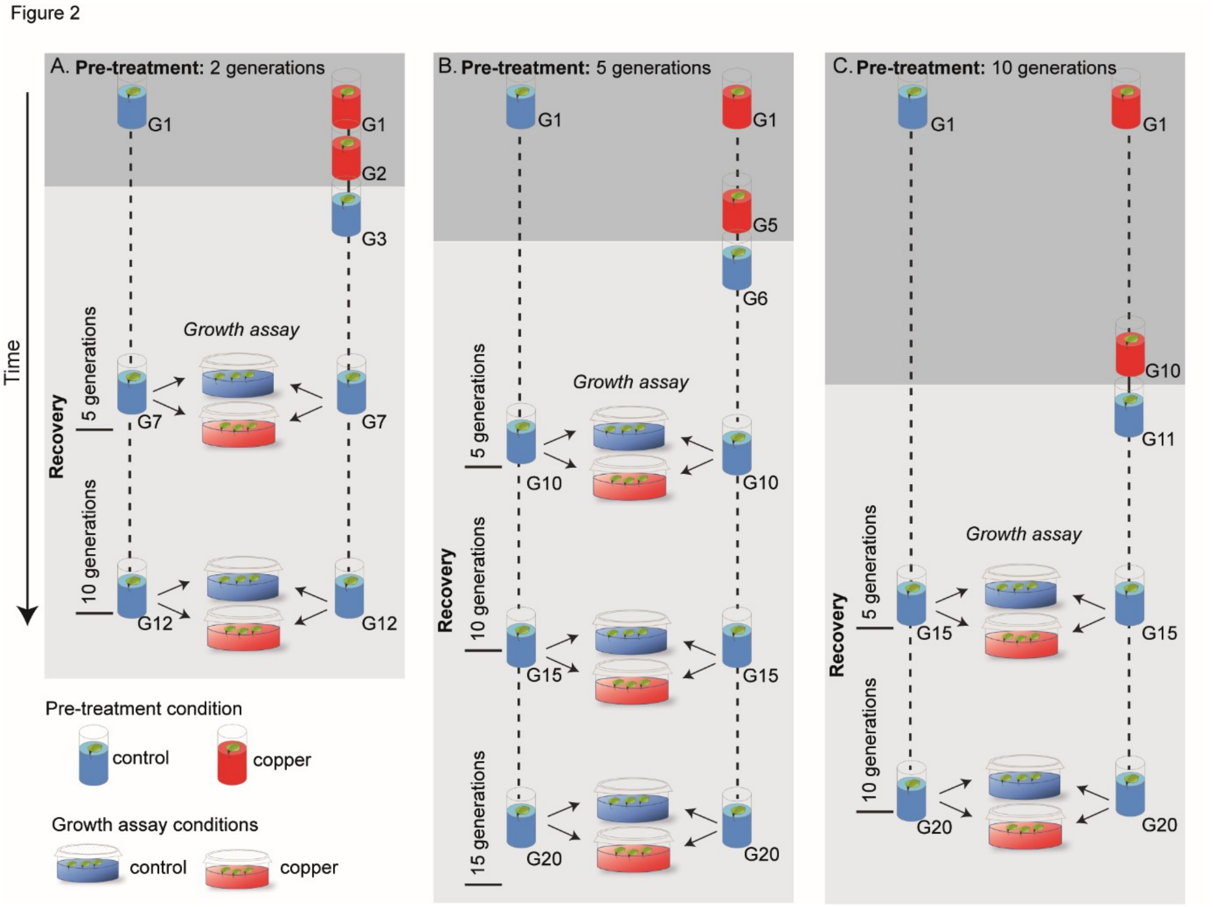


Figure S1. Experimental scheme for transgenerational experiments with two generations (A), 5 generations (B) or 10 generations (C) pre-treatment, and 5, 10 or 15 generations recovery prior performing offspring growth assays across 8 days (2-3 generations). Plants were grown as single descendants during the pre-treatment and recovery phase.

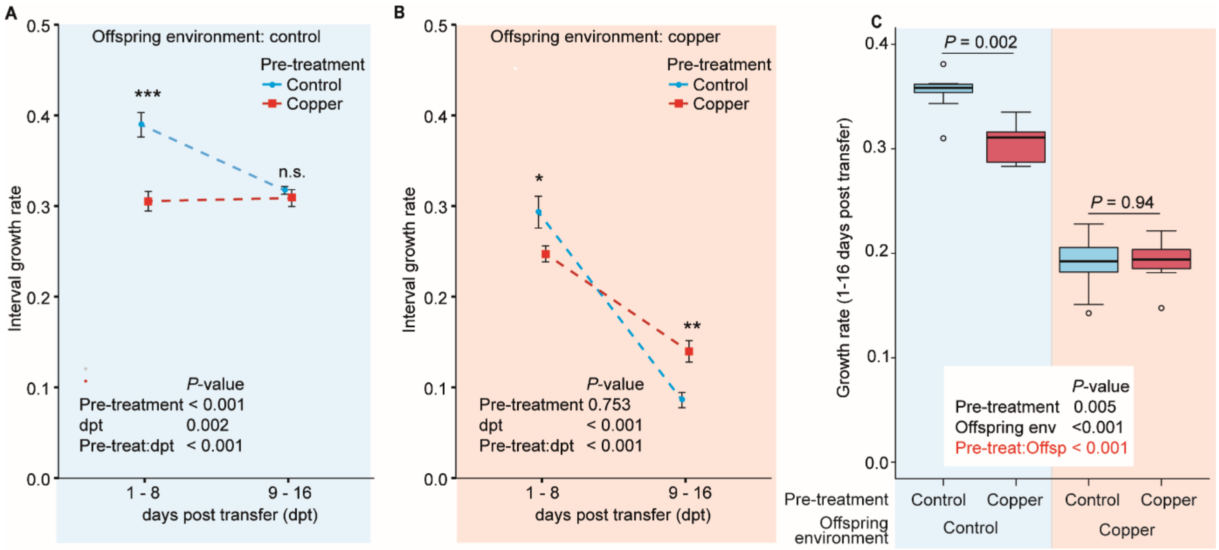


Figure S2. Offspring growth rates of copper and control pre-treated monoclonal *S. polyrhiza* populations across 16 days of growth under control (A) and copper excess (B). Growth rates were assessed in 8-day growth intervals (A and B) as well as across 16 days (C). Asterisks depict *P*-values of Kruskal-Wallis rank sum tests comparing growth rates among pre-treatments. *P*-values of two-way ANOVAs are displayed at the bottom of each panel. (**P* < 0.05, ***P* < 0.01, ****P* < 0.001; n.s. = non-significant). The significant interaction of pre-treatment and offspring environment (highlighted in red) refers to alterations in plant resistance (C). N = 9-10.

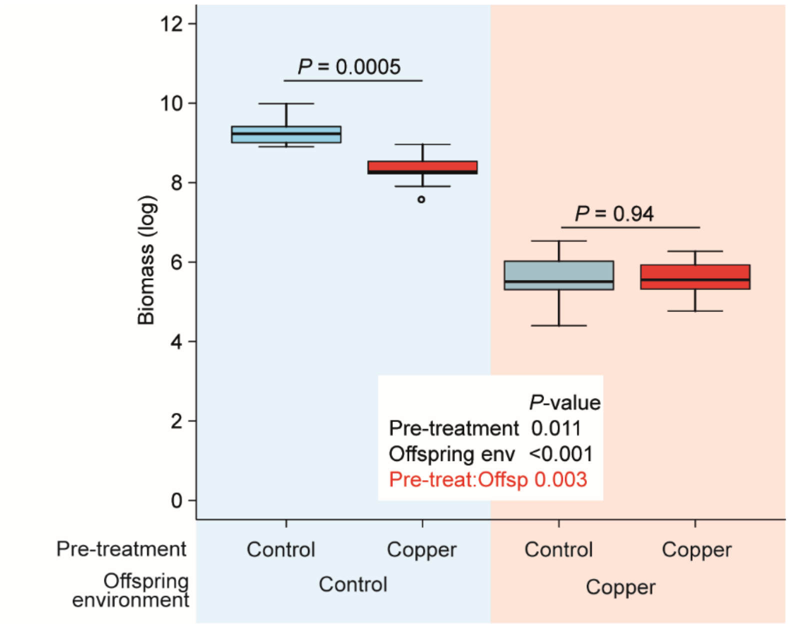


Figure S3. Offspring biomass accumulation (log-transformed) of control and copper pre-treated monoclonal *S. polyrhiza* populations across 16 days of growth. *P*-values of Kruskal-Wallis rank sum tests and two-way ANOVA are shown. The significant interaction of pre-treatment and offspring environment (highlighted in red) refers to alterations in plant resistance. N = 9-10.

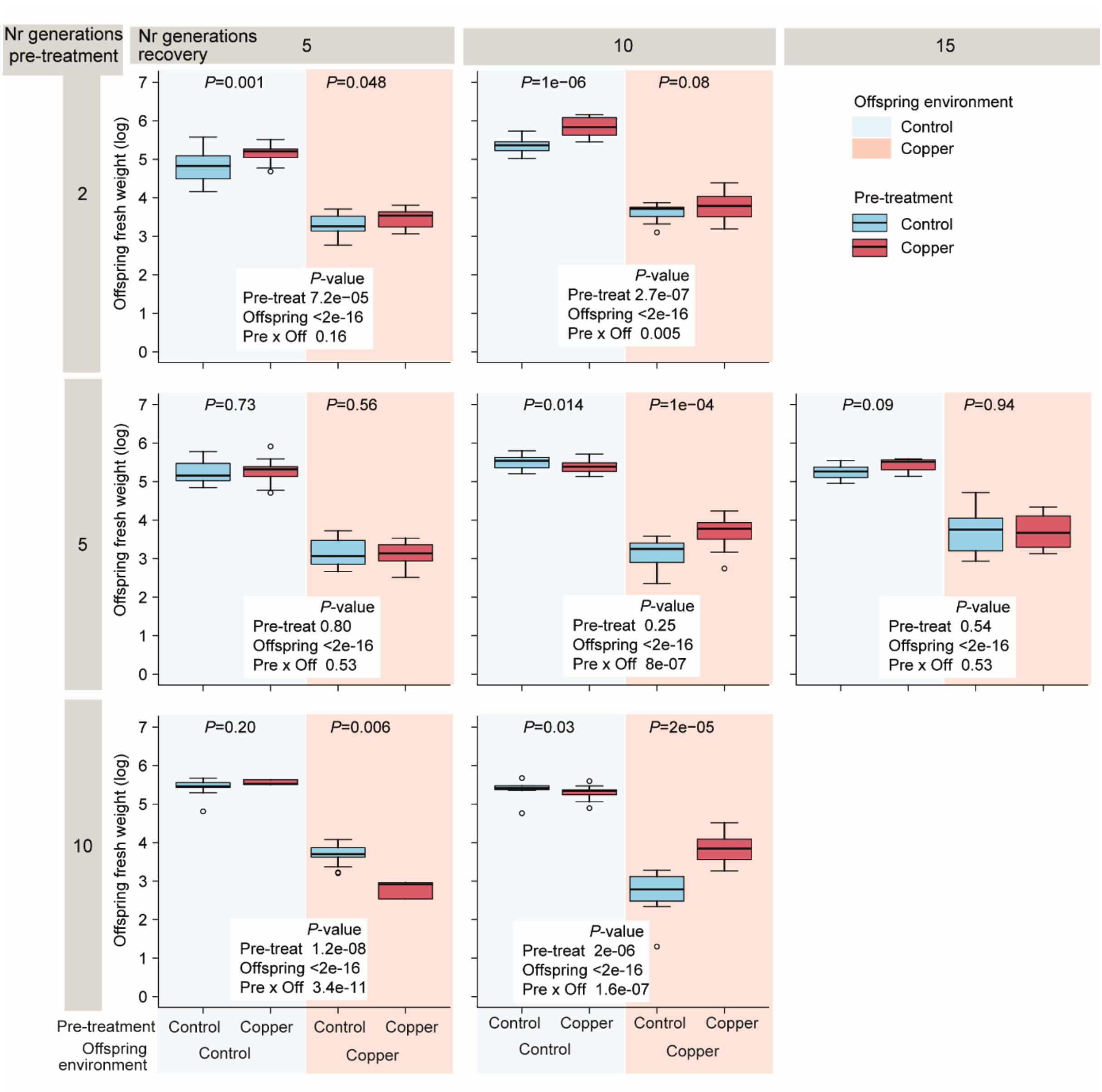


Figure S4. Accumulation of offspring fresh weight (log-transformed) after 8 days of growth in the presence and absence of recurring copper excess at different durations of the pre-treatment and recovery phase. *P*-values refer to two-way ANOVAs and Kruskal-Wallis rank sum tests. N = 3-25.

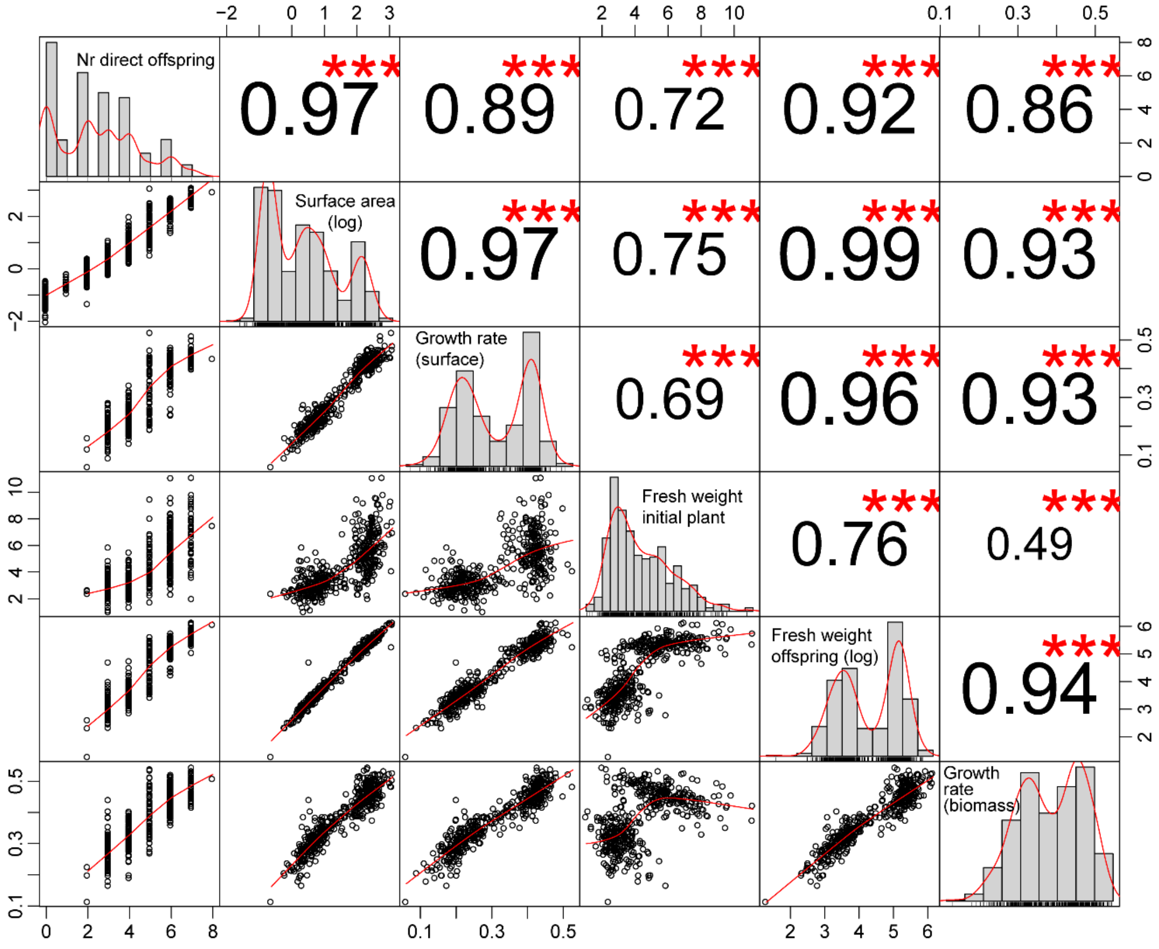


Figure S5. Correlation among fitness parameters assessed in the transgenerational experiment. Pearson's product moment correlation coefficients and *P*-value of Pearson's correlation tests are shown as asterisks (* $P < 0.05$, ** $P < 0.01$, *** $P < 0.001$). Surface area and fresh weight of offspring were log-transformed prior analysis.

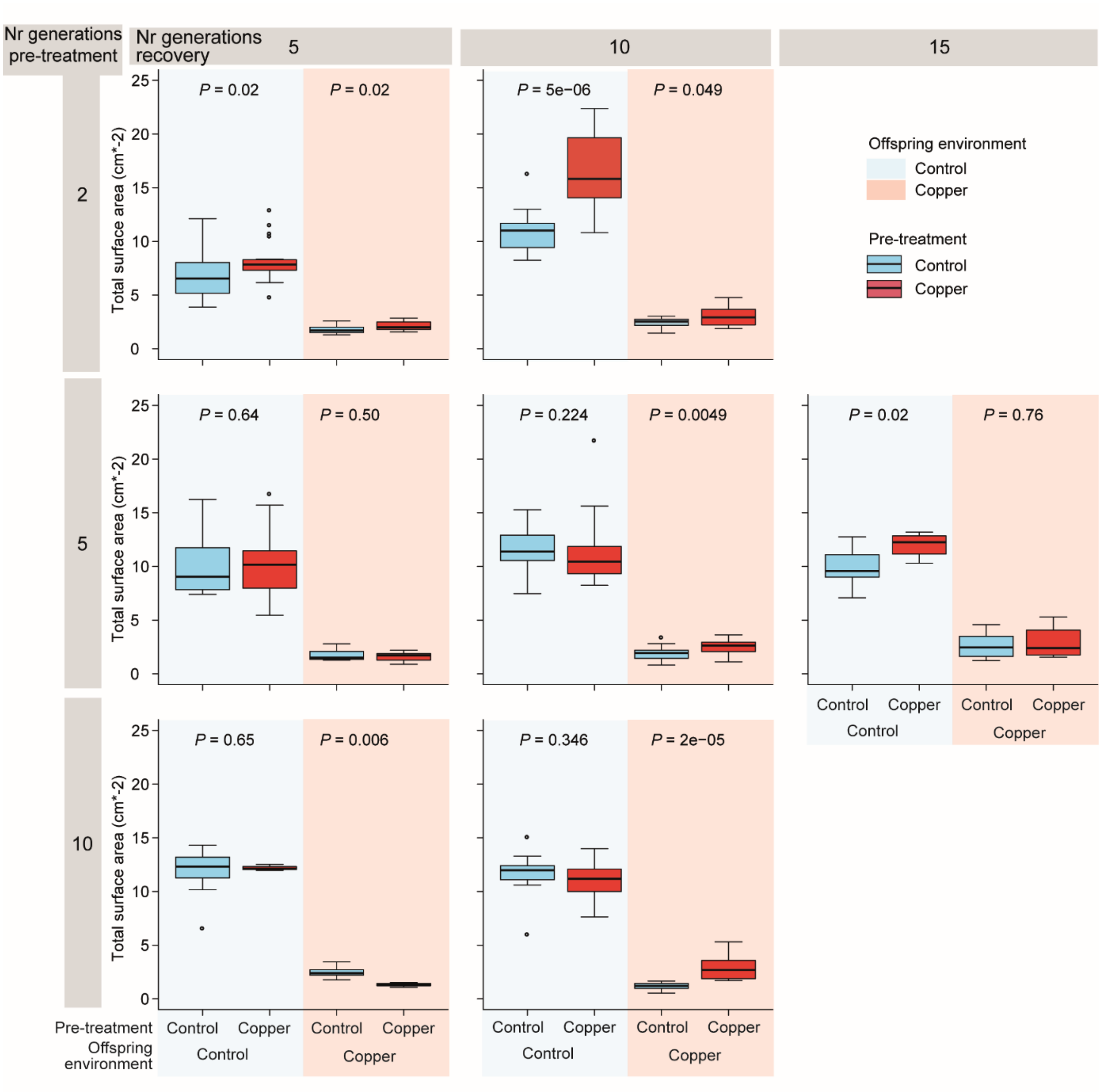


Figure S6. Frond surface area of offspring after 8 days of growth in the presence and absence of recurring copper excess at different durations of the pre-treatment and recovery phase. *P*-values refer to Kruskal-Wallis rank sum tests. N = 3-25.

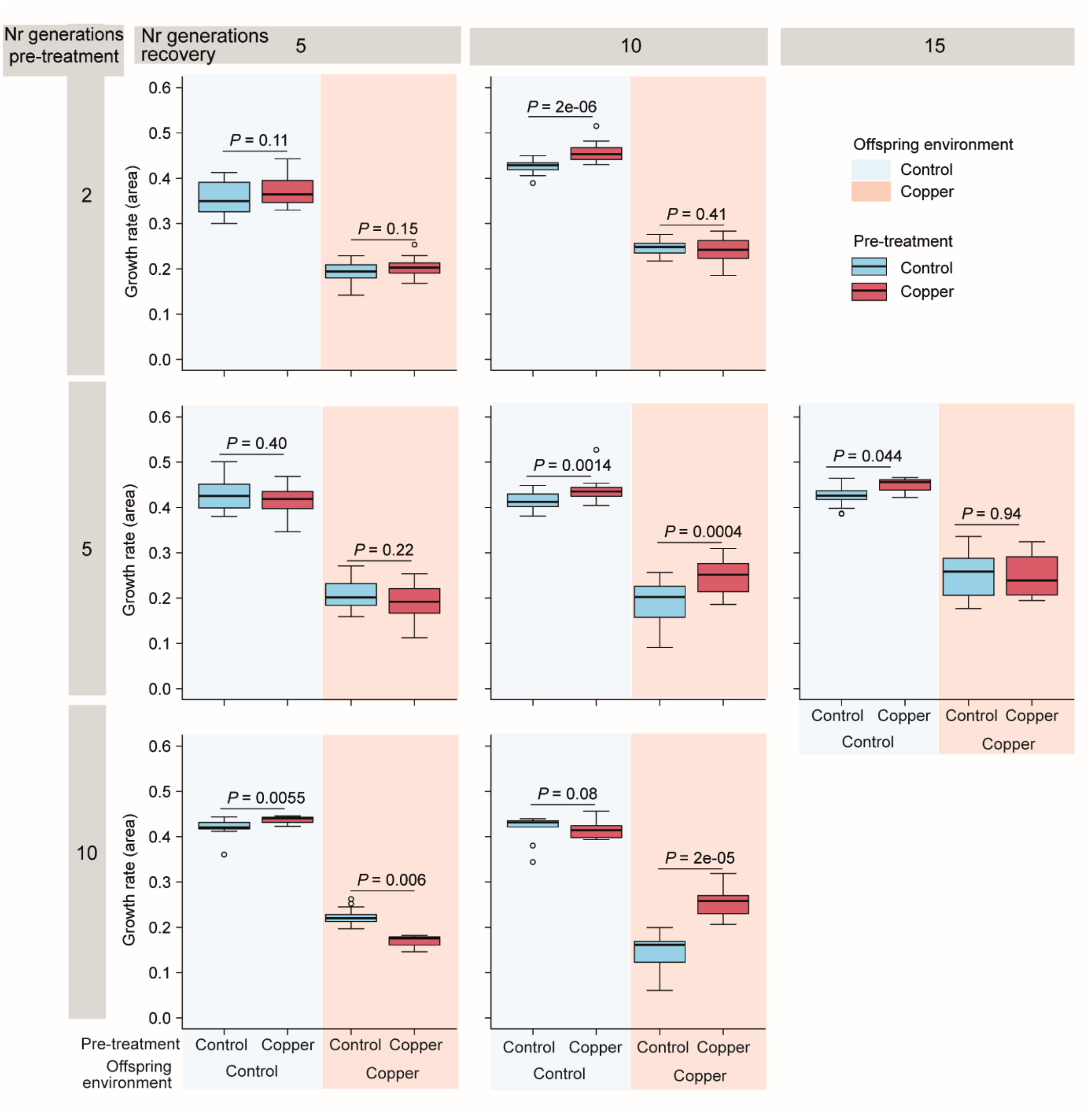


Figure S7. Area-based growth rates of offspring across 8 days of growth in the presence and absence of recurring copper excess at different durations of the pre-treatment and recovery phase. *P*-values refer to Kruskal-Wallis rank sum tests. N = 3-25.

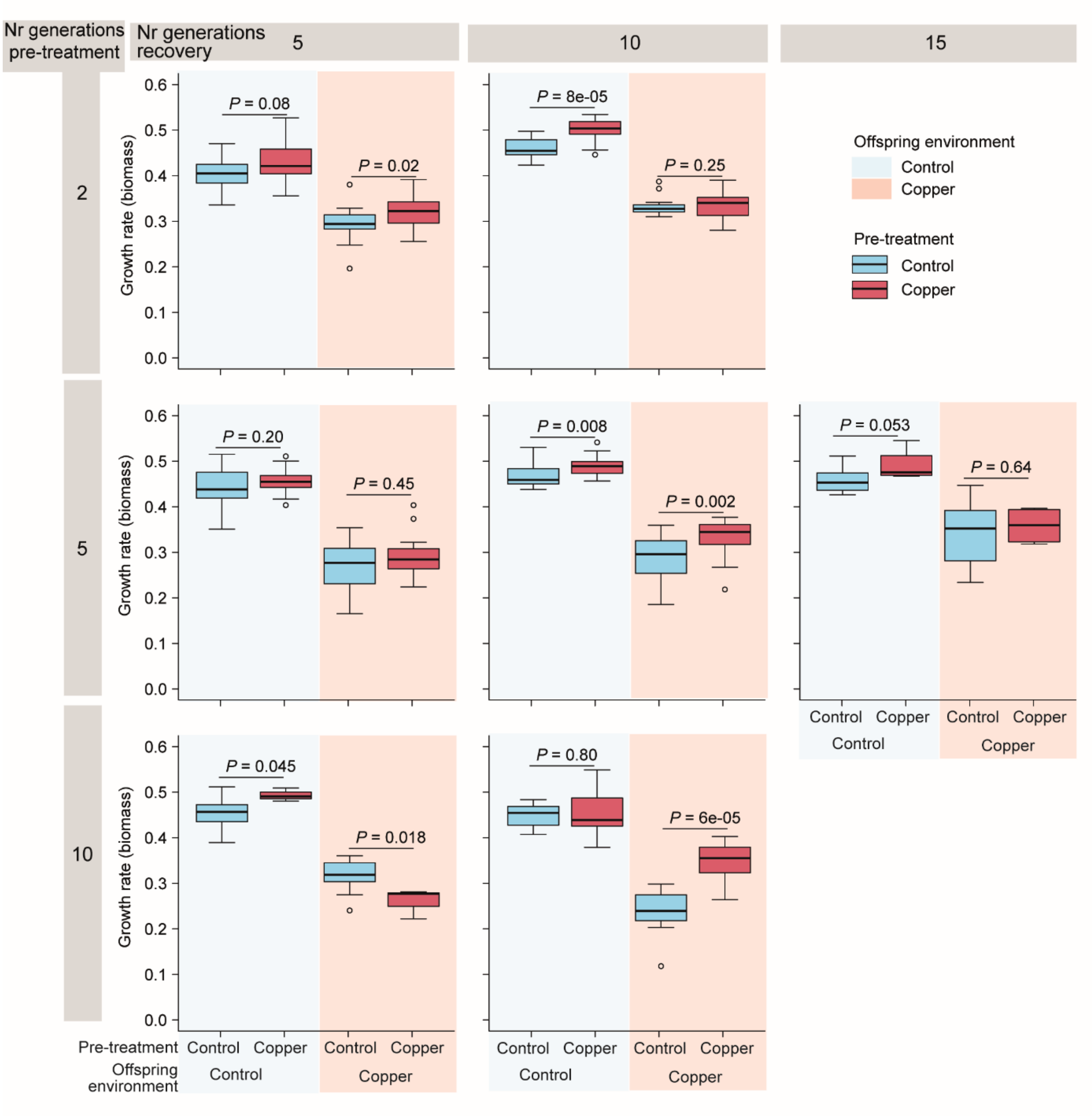


Figure S8. Fresh weight-based growth rates of offspring across 8 days of growth in the presence and absence of recurring copper excess at different durations of the pre-treatment and recovery phase. *P*-values refer to Kruskal-Wallis rank sum tests. N = 3-25.

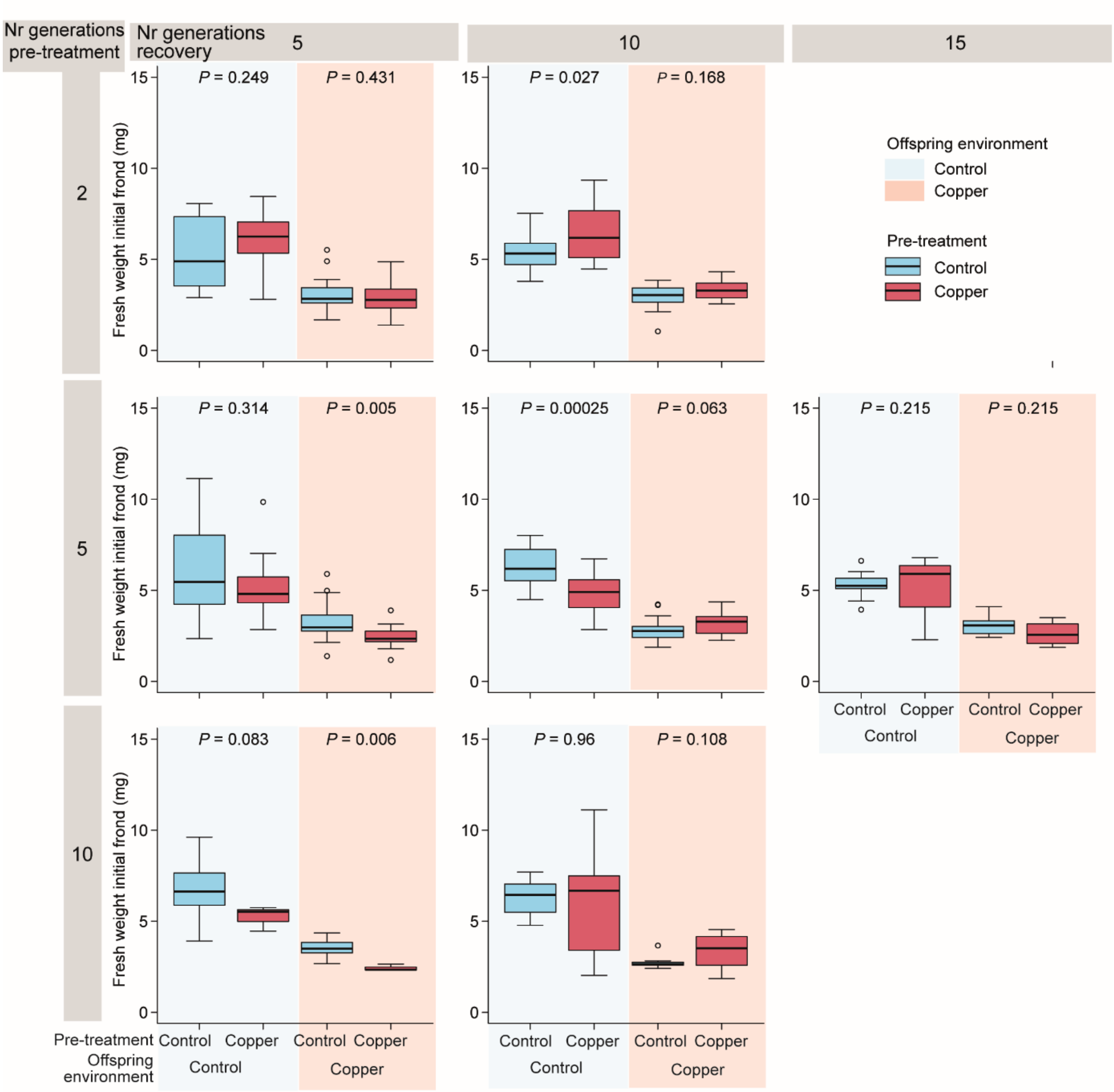


Figure S9. Fresh weight of the initial frond (used to initiate offspring growth assays) after 8 days of growth in the presence and absence of recurring copper excess at different durations of the pre-treatment and recovery phase. *P*-values refer to Kruskal-Wallis rank sum tests. N = 3-25.

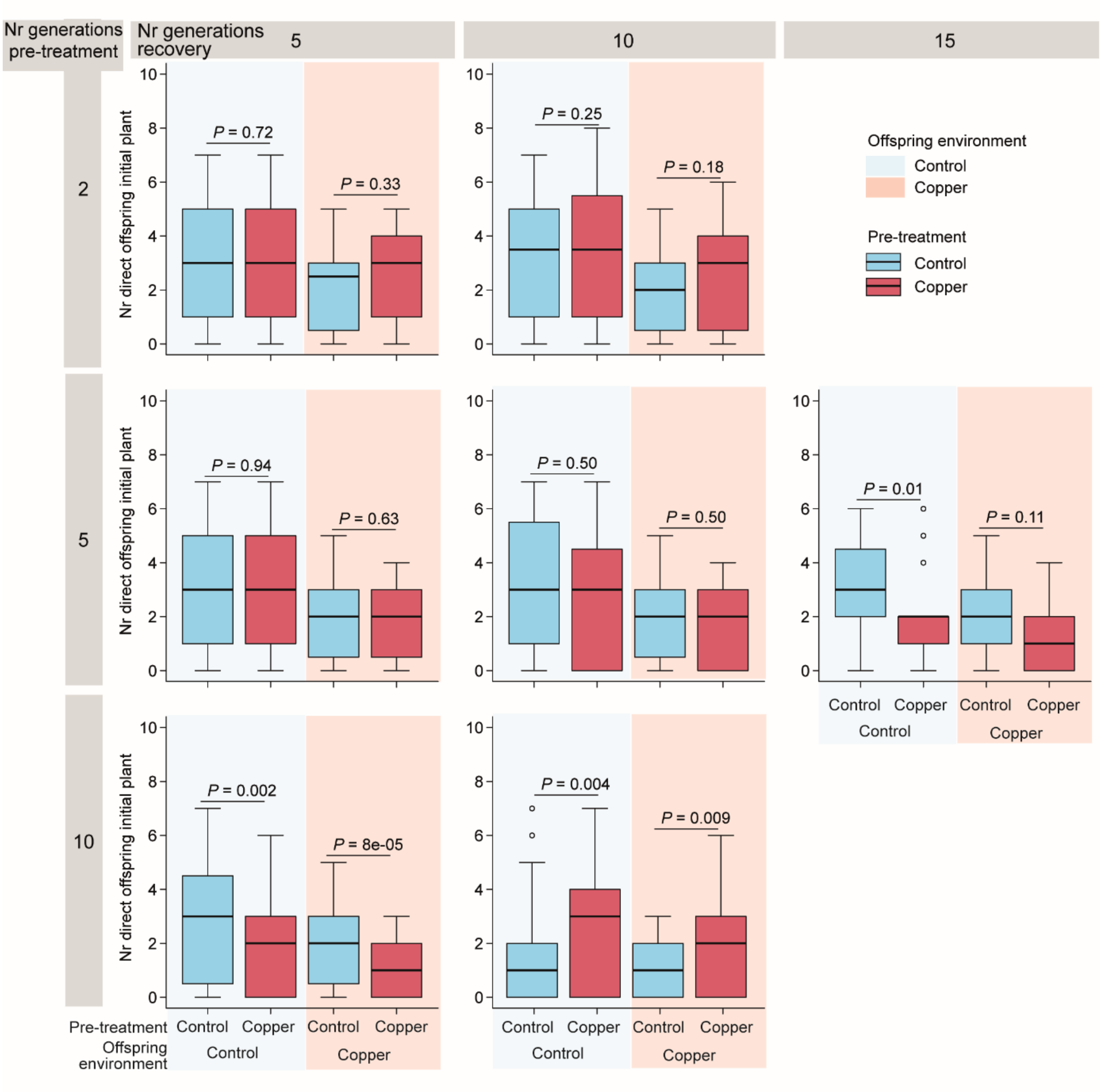


Figure S10. Number of daughters emerging from the initial frond (used to initiate offspring fitness assays) during 8 days of growth in the presence and absence of recurring copper excess at different durations of the pre-treatment and recovery phase. *P*-values refer to Kruskal-Wallis rank sum tests. N = 3-25.

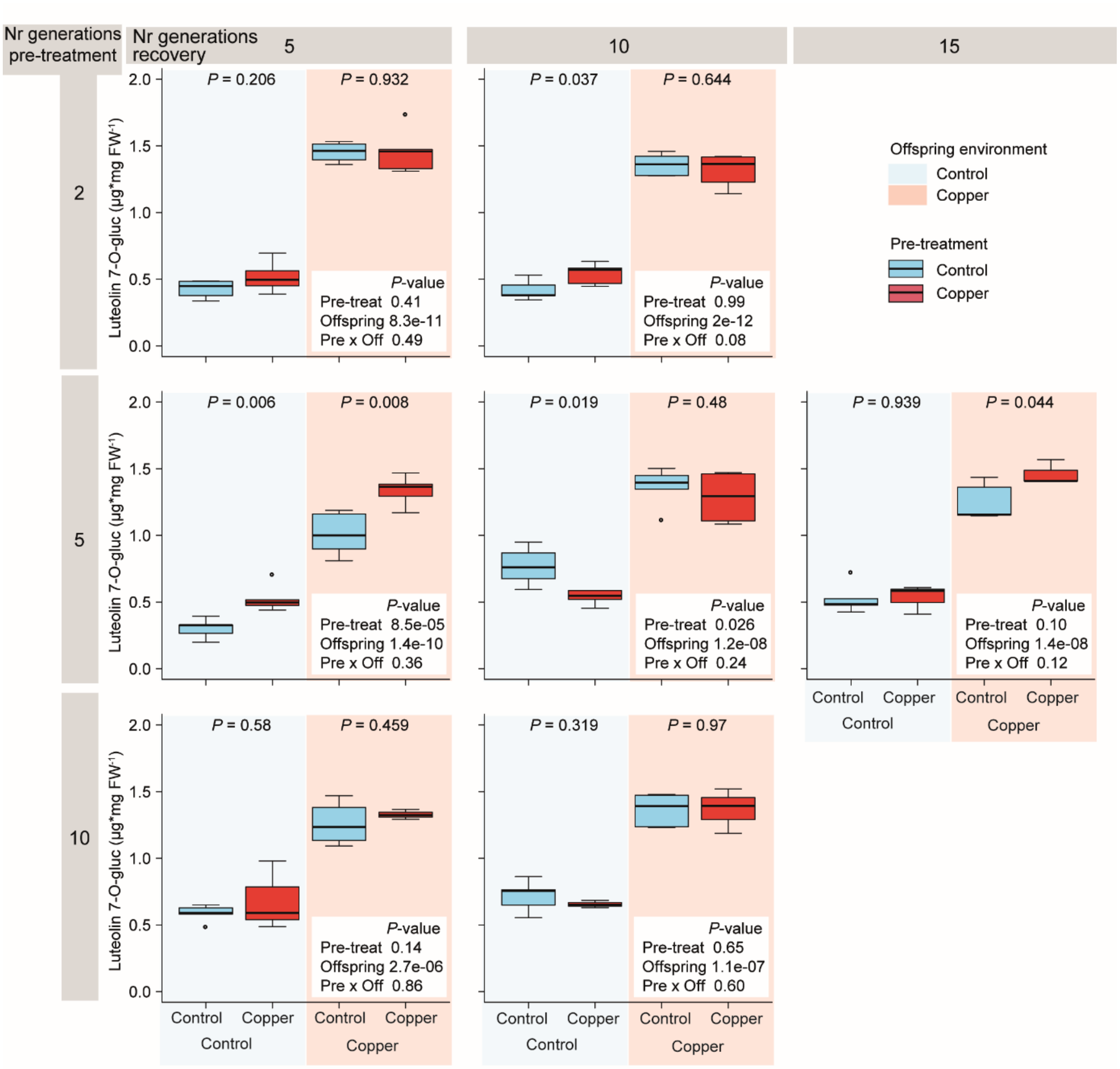


Figure S11. Concentration of luteolin 7-O-glucoside in offspring after 8 days of growth in the presence and absence of recurring copper excess at different durations of the pre-treatment and recovery phase. *P*-values of Student's *t*-tests and two-way ANOVAs are shown. N = 3-6. FW = fresh weight.

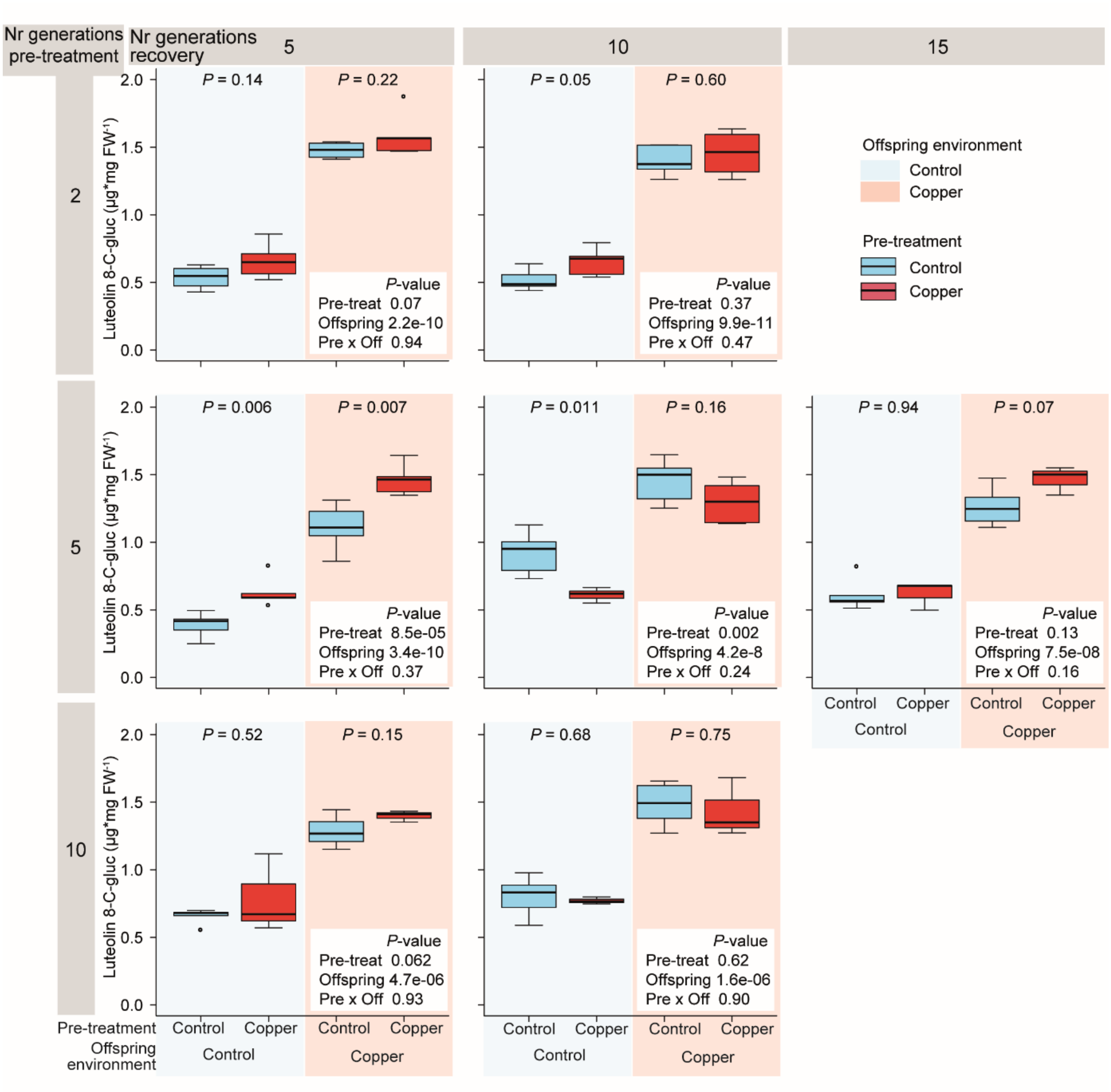


Figure S12. Concentration of luteolin 8-C-glucoside in offspring after 8 days of growth in the presence and absence of recurring copper excess at different durations of the pre-treatment and recovery phase. *P*-values of Student's *t*-tests and two-way ANOVAs are shown. N = 3-6. FW = fresh weight.

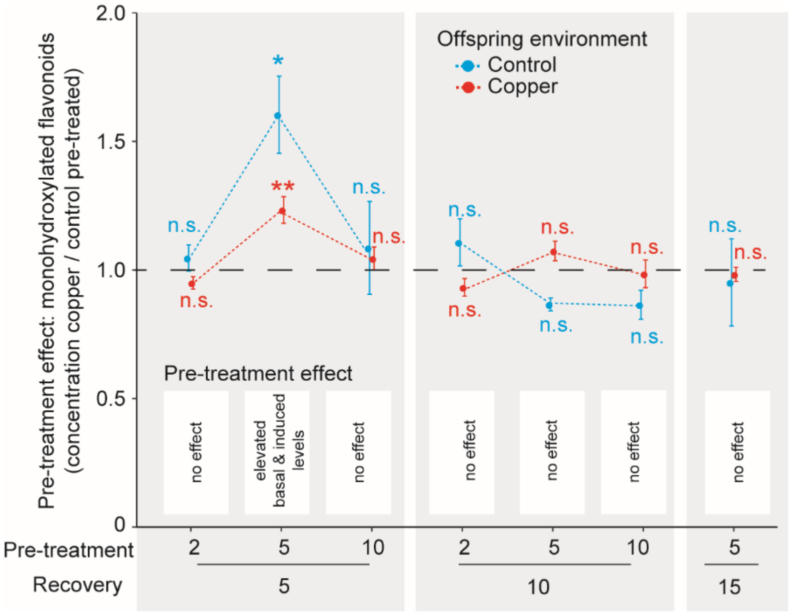


Figure S13. Concentration of monohydroxylated B-ring-substituted flavonoids (sum of apigenin 7-O- and apigenin 8-C-glucosides) in offspring after 8 days of growth in the presence and absence of copper excess at different durations of the pre-treatment and recovery phase. *P*-values of Student's *t*-tests comparing the concentrations of copper and control pre-treated offspring grown under control or copper excess are shown. N = 3-6. FW = fresh weight.

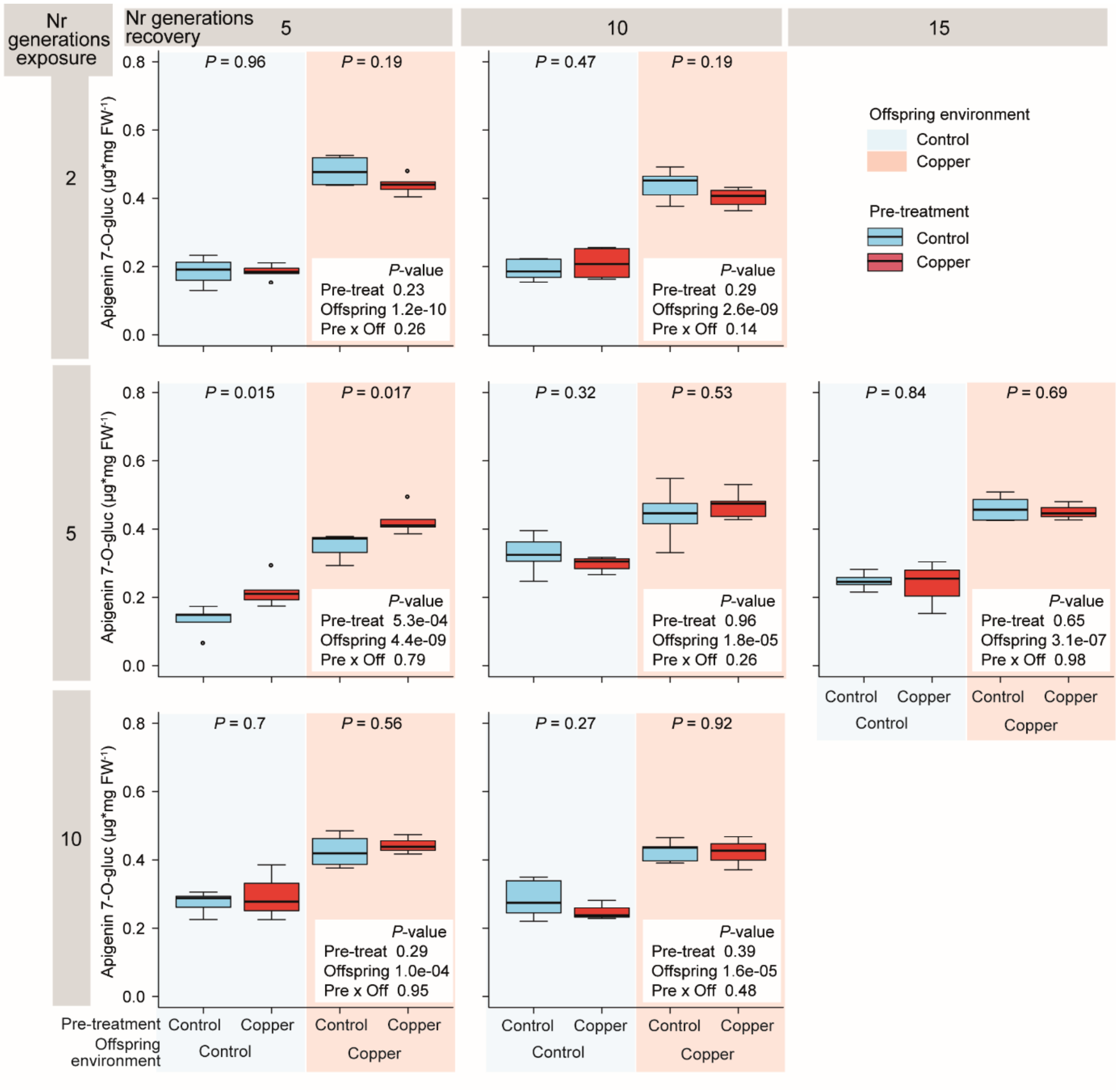


Figure S14. Concentration of apigenin 7-O-glucoside in offspring after 8 days of growth in the presence and absence of recurring copper excess at different durations of the pre-treatment and recovery phase. *P*-values of Student's *t*-tests and two-way ANOVAs are shown. N = 3-6. FW = fresh weight.

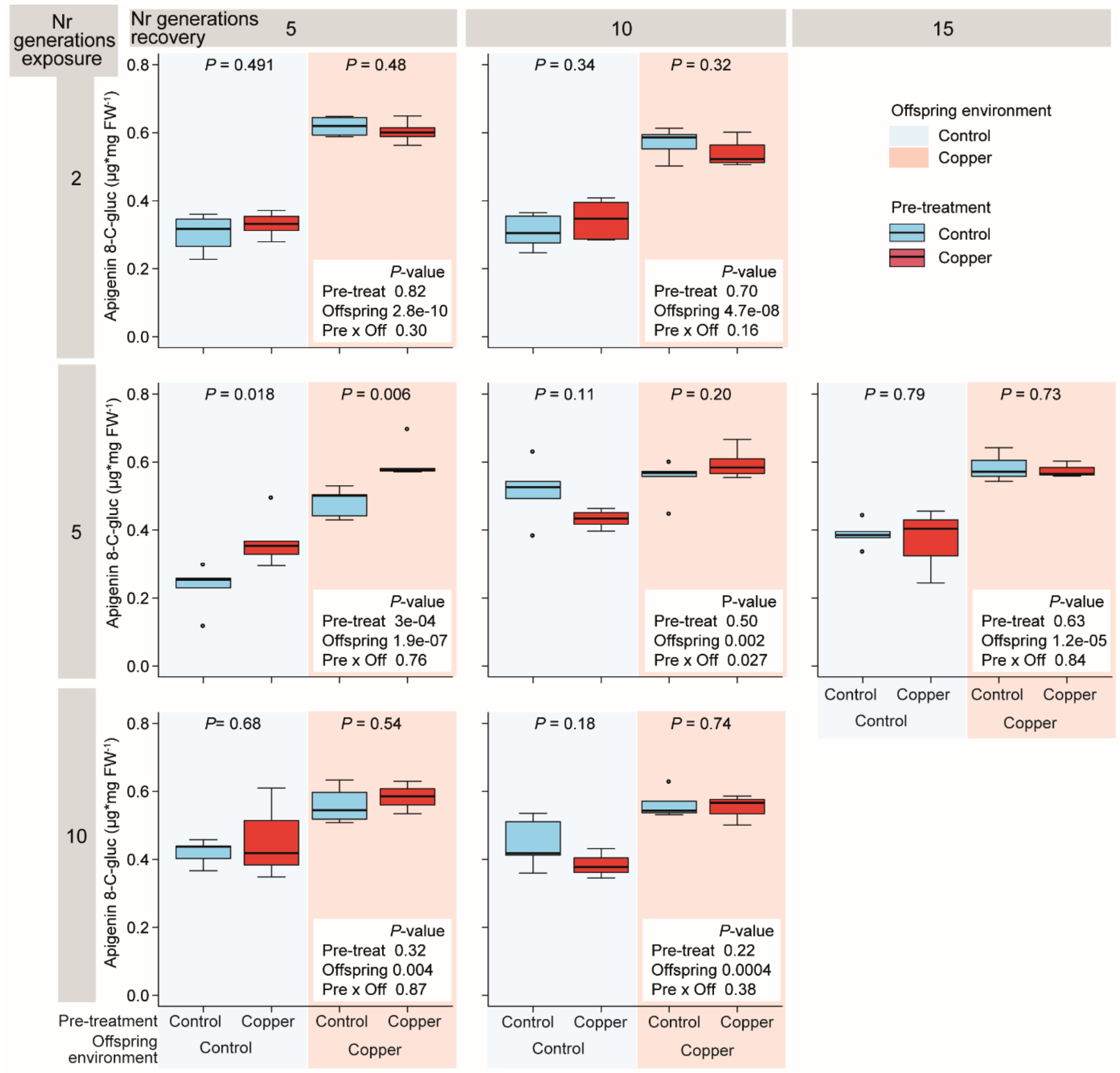


Figure S15. Concentration of apigenin 8-C-glucoside in offspring after 8 days of growth in the presence and absence of recurring copper excess at different durations of the pre-treatment and recovery phase. *P*-values of Student's *t*-tests and two-way ANOVAs are shown. N = 3-6. FW = fresh weight.

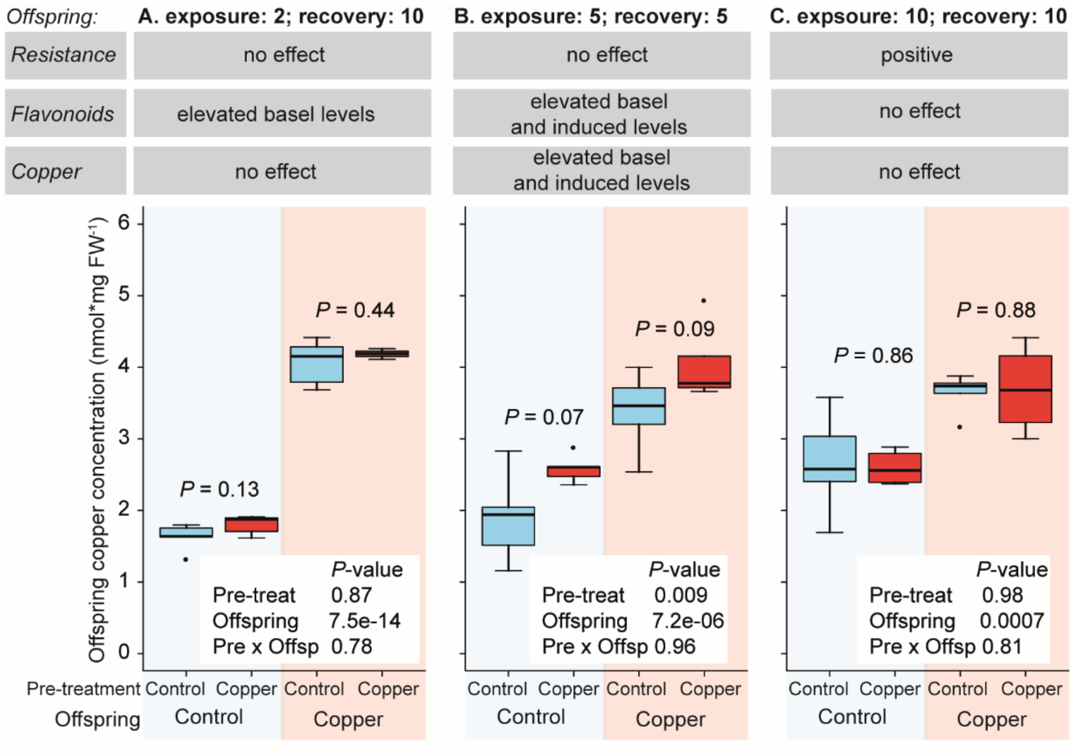


Figure S16. Copper concentration of copper and control pre-treated offspring under different pre-treatment and recovery phase durations (A-C). Offspring copper concentration was measured after 8 days of growth in the presence and absence of copper excess. *P*-values of Student's *t*-tests and two-way ANOVAs are shown. Offspring resistance and flavonoid accumulation patterns are display above the panel and refer to results of figure 3 and 4. N = 4-5.

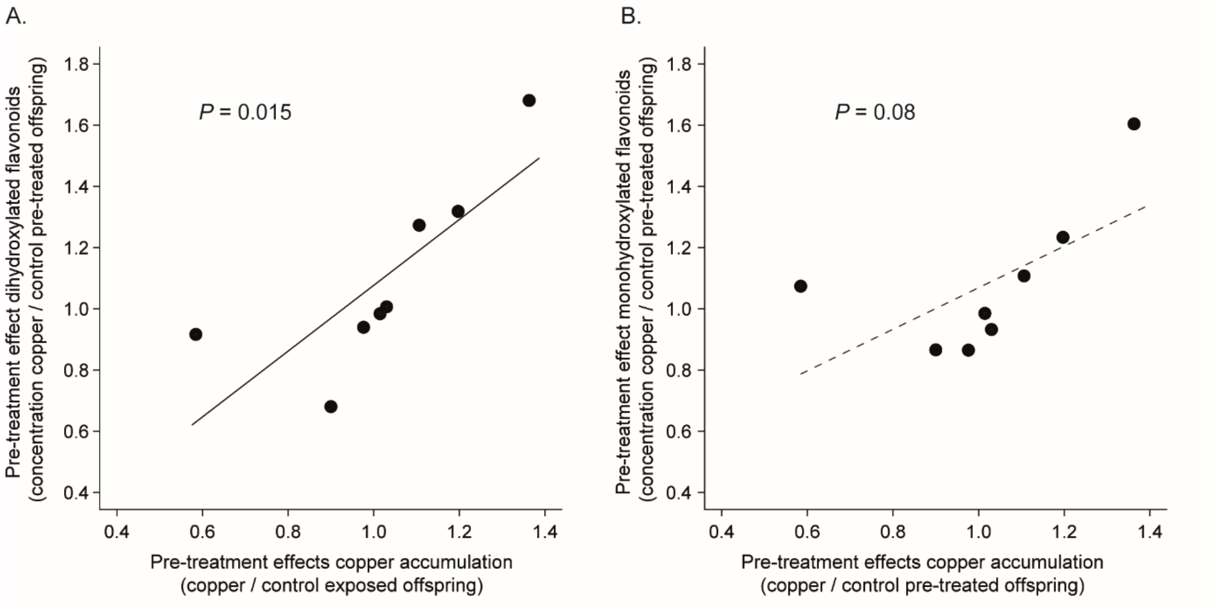


Figure S17. Pre-treatment effects (concentration of copper to control pre-treated offspring) among offspring copper and dihydroxylated (A) and monohydroxylated (B) B-ring substituted flavonoid accumulation. *P*-values of linear models are shown. Each data point displays the mean value of a pre-treatment and recovery combination and offspring environment. N = 8.
